# Blocking glutamine transport normalizes lymphatic vessels in hypoxic environments by attenuating glycolysis

**DOI:** 10.64898/2025.12.18.695240

**Authors:** Ellie Johandes, Eva Hall, Tomas Harbut, Kaden Priebe, Margaret A. Schwarz, Donny Hanjaya-Putra

## Abstract

Dysfunctional lymphangiogenesis is a component of several diseases, including secondary lymphedema, allergic asthma, and solid malignancies. These vessels are ineffective at draining interstitial fluid, resulting in secondary complications such as edema, increased inflammation, slowed wound healing, and, for cancer patients, increased risk of metastasis. Hypoxic microenvironments drive dysfunctional vessel growth by upregulating vascular endothelial growth factor receptors, migratory signaling pathways, and glycolytic metabolism. Despite the potential benefits of normalizing lymphatic vasculature, most treatments for vascular normalization focus on blood vasculature, ignoring the unique properties of lymphatic endothelial cells (LECs). Moreover, previous targets for vascular normalization center around vascular endothelial growth factors, which risk adverse side effects. To address these issues, we approached lymphatic normalization from a metabolic perspective. In this study, we investigated the impact of glutamine availability on factors critical to lymphangiogenesis, including glycolysis, cell proliferation, and migration. We found that increasing the concentration of glutamine in media results in increased lactate production and the expression of glycolytic genes *HK2*, *GLUT1*, and *GLUT3* under hypoxia. The presence of glutamine also encouraged LEC proliferation, while blocking glutamine transport reduced lactate production, *HK2* expression, and slowed collective LEC migration. In a 2D vessel formation assay, we found that glutamine increased vessel formation in normoxic conditions but lowered vessel connectivity in hypoxic conditions, reflecting the dysfunction seen in hypoxic diseases. However, attenuating glycolysis by blocking glutamine transport caused LECs to form longer, interconnected vascular networks. This study reveals that glutamine availability can modulate LEC glycolysis, and therefore lymphangiogenesis, in a hypoxia-dependent manner.

## Introduction

The lymphatic system is responsible for fluid balance, waste removal, and immune cell trafficking. In healthy tissues, new lymphatic vessels will sprout in response to tissue damage, driven by vascular endothelial growth factors (VEGFs), inflammation, and hypoxia (Zampell et al., 2012). These new vessels speed healing by draining away inflammatory cytokines and resolving edema. In mice, increased lymphangiogenesis has been shown to improve cutaneous wound healing, reduce fibrosis post myocardial infarction, and protect against neurological damage from stroke (Boisserand et al., 2024; Pollack et al., 2025; Vieira et al., 2018). If inflammation and hypoxia persist, however, lymphangiogenesis can become dysfunctional. Rather than forming organized, stable vasculature, new vessels are disorganized and leaky. Impaired fluid drainage can result in edema, increased inflammation, hypoxia, fibrosis, and slowed wound healing (Jiang et al., 2022). Dysfunctional lymphatics exacerbate chronic conditions such as secondary lymphedema and allergic asthma, as well as provide convenient pathways for metastasis in the solid tumor microenvironment (Gomez Medellin et al., 2024; Ji et al., 2023; Jiang et al., 2022; Qiu et al., 2024).

Despite the importance of lymphatic vasculature, much of our current knowledge of vascular dysfunction and normalization comes from studies on blood endothelial cells (BECs) (Cao et al., 2019; Choi and Jung, 2023; Yang et al., 2022). While lymphatic endothelial cells (LECs) share many similarities to BECs, including their reliance on glycolysis for energy and surface expression of CD31 (PECAM-1), the lymphatic system’s role in lymph transport gives them a distinct identity (Culic et al., 1997; De Bock et al., 2013; De Bock et al., 2013; Muller et al., 1989; Yu et al., 2017). Unique to LECs is their expression of transcription factor PROX1 and glycoprotein Podoplanin, which are responsible for the development of lymphatic vasculature and maintenance of lymphatic identity (Cha et al., 2018; Hong et al., 2002; Wang et al., 2023). LYVE1, a hyaluronan receptor, acts as a docking site for dendritic cells and assists in the migration of macrophages into the lymphatic system (Jackson, 2019). While both BECs and LECs can form vessels in response to vascular endothelial growth factor (VEGF)-A, VEGF-C is the main driver of lymphangiogenesis. Furthermore, compared to other endothelial cells (ECs), LECs express more genes associated with vesicular transport (Lee et al., 2025; Nelson et al., 2007; Podgrabinska et al., 2002). In terms of behavior, LECs have unique patterns of cell-cell adhesion, paracrine signaling, and methods of contact inhibition compared to BECs (Carlantoni et al., 2025; Jeong et al., 2022; Kriehuber et al., 2001; Lee et al., 2023). As it should not be assumed that previous techniques for vascular normalization applied to BECs will work as effectively for LECs (Saha et al., 2023), this study seeks to find new methods to curb hypoxia-driven lymphangiogenesis.

Hypoxia spurs vessel growth through the stabilization of hypoxia-inducible factors (HIFs) (Wang et al., 1995). In endothelial cells (ECs), HIFs spur (lymph)angiogenesis by upregulating migratory signaling pathways, increasing the production of VEGF and its receptors, and shifting cell metabolism towards increased glycolysis (Abaci et al., 2010; Liu et al., 2025). As such, blocking VEGF receptors are a main target to control vessel growth. However, anti-angiogenic treatments affect both healthy and dysfunctional vessels, resulting in adverse side effects ranging from hypertension to intestinal bleeding (Choi and Jung, 2023; Goel et al., 2011). Therefore, a treatment which specifically targets diseased vessels is needed.

ECs undergo large metabolic changes between quiescent and angiogenic states. Recent work by Durot *et al*. (2025) showed that proliferating ECs rely on glycolysis but shift to fatty acid oxidation when they become quiescent. Furthermore, glycolysis directly powers EC migration and vessel sprouting. This can result in endothelial barrier dysfunction in the form of increased permeability (Gallemit et al., 2021; Yang et al., 2022). As such, researchers have turned to limiting endothelial glycolysis as a mechanism to normalize vasculature (Li et al., 2019). Previous works in BECs have demonstrated that inhibiting glycolytic enzyme PFKFB3 can inhibit vascular sprouting *in vitro* and normalize tumor vasculature for improved drug delivery in mice (Abdali et al., 2021; Cantelmo et al., 2016). Success has also been found by co-inhibiting pyruvate dehydrogenase and glutaminase-1 (Schoonjans et al., 2020). In LECs, reducing the translation of glycolytic enzyme hexokinase II (HK2) with miR-484 reduced pathological lymphangiogenesis in mouse corneas (Zhang et al., 2025). However, given the importance of glycolysis in active and quiescent ECs, over-inhibition can result in cell death (De Bock et al., 2013). Rather than targeting glycolysis directly, we investigate whether it can be modulated through other metabolic pathways, in this case glutamine metabolism.

Glutamine is the most abundant amino acid in the blood. Its metabolites contribute to multiple cellular pathways in LECs and BECs, including nucleotide synthesis, lipid synthesis, glutaminolysis, and redox homeostasis (Mirveis et al., 2023; Teuwen et al., 2019; Yoo et al., 2020b). Although a direct link between glutamine availability and glycolysis has not previously been shown in ECs, glutamine is also highly consumed in other glycolysis-reliant cells, including lymphocytes, stem cells, and some cancers (Carr et al., 2010; Loopmans et al., 2024; Pérez-Escuredo et al., 2016; Tohyama et al., 2016; van Geldermalsen et al., 2016; Zhang et al., 2020). As in glycolysis, the transcription of genes associated with glutamine metabolism are also increased in hypoxic conditions via HIF signaling, marking it as a promising target for hypoxic LECs (Sun and Denko, 2014; Xiang et al., 2019).

In this study, we seek to determine whether controlling glutamine availability can influence glycolysis, and therefore lymphangiogenesis, in normoxic and hypoxic LECs. First, we clarify the relationship between glutamine availability and glycolysis by measuring lactate production along with the expression of glycolytic genes hexokinase II (*HK2*), glucose transporter 1 (*GLUT1*) and glucose transporter 3 (*GLUT3*) over increasing concentrations of glutamine. We then repeat these experiments while pharmacologically inhibiting glutamine transport. Then, we assess glutamine metabolism as a new target for lymphatic vessel normalization by evaluating its impact on LEC proliferation, migration, and vessel formation *in vitro*. Our results indicate that hypoxic conditions increase the production of glutamate by LECs. Increasing glutamine concentration in the media reveals a hypoxia-specific increase in *GLUT1* and *GLUT3* expression alongside a non-significant increase in *HK2*. We also found that glutamine increases LEC proliferation and migration, while its effect on vessel formation is oxygen-dependent. As such, this study offers preliminary evidence that glutamine availability can influence LEC glycolysis, making it a future target for curbing dysfunctional lymphangiogenesis in chronic, hypoxic diseases.

## 2. Results

### 2.1. Hypoxia and glutamine availability increase glutamine utilization by LECs

Hypoxia increases glutamine uptake in skeletal stem cells, T-cells, and various cancers (Sun and Denko, 2014; Yoo et al., 2020a; Loopmans et al., 2024; Wik et al., 2022). To observe whether this trend continues in lymphatic endothelial cells, LECs were cultured for 24 hours in a hypoxic incubator set to 5% CO_2_, 94% N_2_, and 1% oxygen by volume. Oxygen levels in the media were measured using a Presens optical oxygen sensor to ensure a final oxygen concentration of < 3%. By t = 24 hours, oxygen levels in the media were measured at 2.45% ± 0.081 **(Fig 1 a-b)**. To further confirm the generation of a hypoxic environment, LECs cultured for 24 hours in normoxia or hypoxia were immunostained for HIF1A **(Fig 1 c)**. In addition to a visual increase in HIF1A, the fluorescent intensity of HIF1A in the nuclei significantly increased **(Fig 1 d)**. To observe whether hypoxia increases glutamine use by LECs, we measured their production of glutamate, which is produced when glutamine is broken down by glutaminase. LECs were serum-starved for approximately 16 hours in glutamine-free DMEM before their media was replaced with glutamine-free DMEM containing Promocell MV2 supplement and 0 - 20 mM of L-glutamine. The cells were then incubated in normoxic or hypoxic conditions at 37 ℃ for 12 hours. Afterwards, media samples were collected and the relative amount of glutamate in the media was measured using a Promega Glutamate-Glo luminescence assay according to manufacturer instructions. In both normoxic and hypoxic conditions, increased glutamine availability increased the amount of glutamate produced. Furthermore, hypoxia significantly increased glutamate production at all levels of glutamine supplementation **(Fig 1e)**. This suggests that both glutamine and oxygen availability influence LEC glutamine utilization. However, it should be noted that glutamate levels can also be increased through the conversion of ɑ-ketoglutarate to glutamate or the reduction of glutamine synthetase activity, which converts glutamate to glutamine, in response to increased glutamine availability (Yelamanchi et al., 2016).

**Figure 1.**
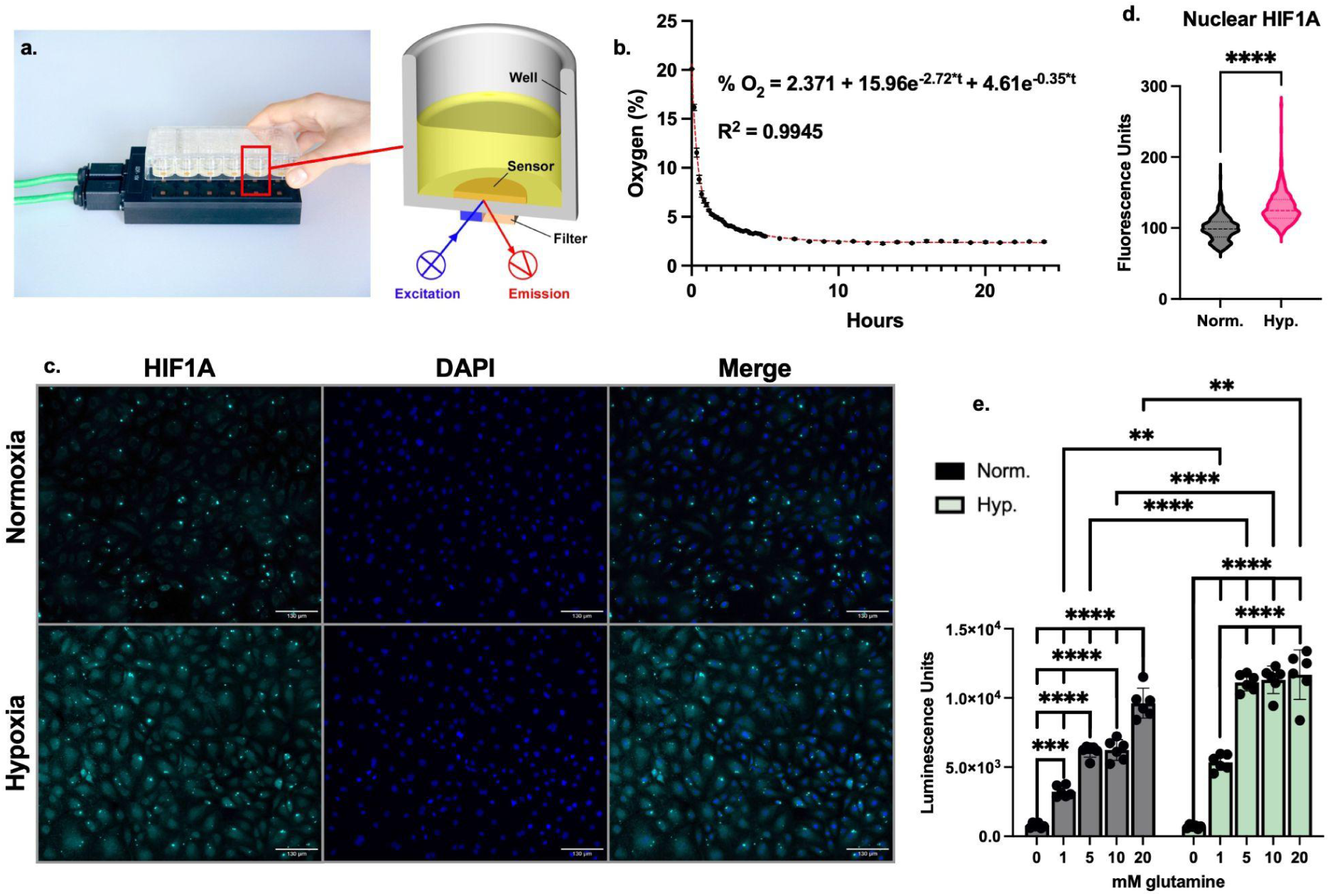
LEC glutamate production increases with both glutamine availability and hypoxia. Percentage of oxygen in cell media was measured using a) a PreSens SDR SensorDish™ reader and OxoHydroDish™ Plate. Images used with permission from PreSens. **b)** Percent oxygen in media over time. Oxygen diffusion from media can be modeled by a two phase decay equation (red trendline), which was calculated in Prism. Each point is this average of n = 3 wells with error bars showing standard deviation. **c)** In addition to monitoring oxygen, the generation of a hypoxic environment was further confirmed by immunostaining for HIF1A (cyan) after 24 hour incubation in normoxic or hypoxic conditions. Cell nuclei were stained with DAPI (blue). Scale bar (white) = 130 μm. **d)** The localization of Hif1α to the nucleus was quantified by measuring the fluorescent intensity of Hif1α in regions overlapping with DAPI. n= 931 nuclei (Norm.); n = 807 nuclei (Hyp.) across 4 wells. Statistical significance determined using Student’s T-test. ****P<0.0001. To investigate changes in glutamine metabolism over changes in oxygen and glutamine availability,glutamate in media was measured by **e)** Promega Glutamate-Glo assay after treatment with 0 - 20 mM of glutamine (n = 6). Glutamate is proportional to luminescence. Statistical significance determined via two-way ANOVA with Tukey’s Test. **P<0.01; ***P<0.001; ****P<0.0001.

### 2.2. Glutamine availability amplifies hypoxia-driven glycolysis

#### 2.2.1 Glutamine supplementation increases glycolytic markers in hypoxic LECs

To determine the relationship between glutamine and glycolytic function, LECs were supplemented with glutamine before incubation in normoxic or hypoxic conditions. Glycolytic activity was assessed by the expression of *HK2*, *GLUT1*, and *GLUT3* mRNA and protein, as well as the presence of lactate in the media. HK2 catalyzes the phosphorylation of glucose, which is the first and rate-limiting step in glycolysis (Roberts and Miyamoto, 2015; Yu et al., 2017). GLUT1 and GLUT3 are the main glucose transporters in endothelial cells (Holman, 2020; Wu and Bai, 2023). Furthermore, the transcription of *HK2*, *GLUT1*, and *GLUT3* are known to be directly upregulated by HIFs in hypoxic conditions (Chen et al., 2001; Riddle et al., 2000; Ryniawec et al., 2022). Finally, lactate is the end-product of glycolysis (Li et al., 2022; Rogatzki et al., 2015). Together, these markers reflect changes in glycolytic activity.

Hypoxia significantly increased the gene expression of *HK2, GLUT1,* and *GLUT3* at all concentrations of glutamine **(Fig 2a-c)**. When examining changes in gene expression due to glutamine supplementation, significant increases to *GLUT1* and *GLUT3* expression were observed under hypoxic conditions. Hypoxic *HK2* expression also increased as glutamine supplementation increased, although this difference was not significant. Gene expression in normoxia was not significantly impacted by glutamine availability with the exception of *GLUT3* at 1 vs 20 mM of glutamine supplementation. These results suggest a positive correlation between glutamine availability and glycolytic gene expression which is enhanced under hypoxic conditions. Interestingly, hypoxic expression of both *GLUT1* and *GLUT3* had an initial, non-significant, drop in expression between 0 and 1 mM treated groups, perhaps indicating a compensatory upregulation of glucose transport when glutamine is absent. This has been previously observed in cancer cells (Schulte et al., 2018).

**Figure 2.**
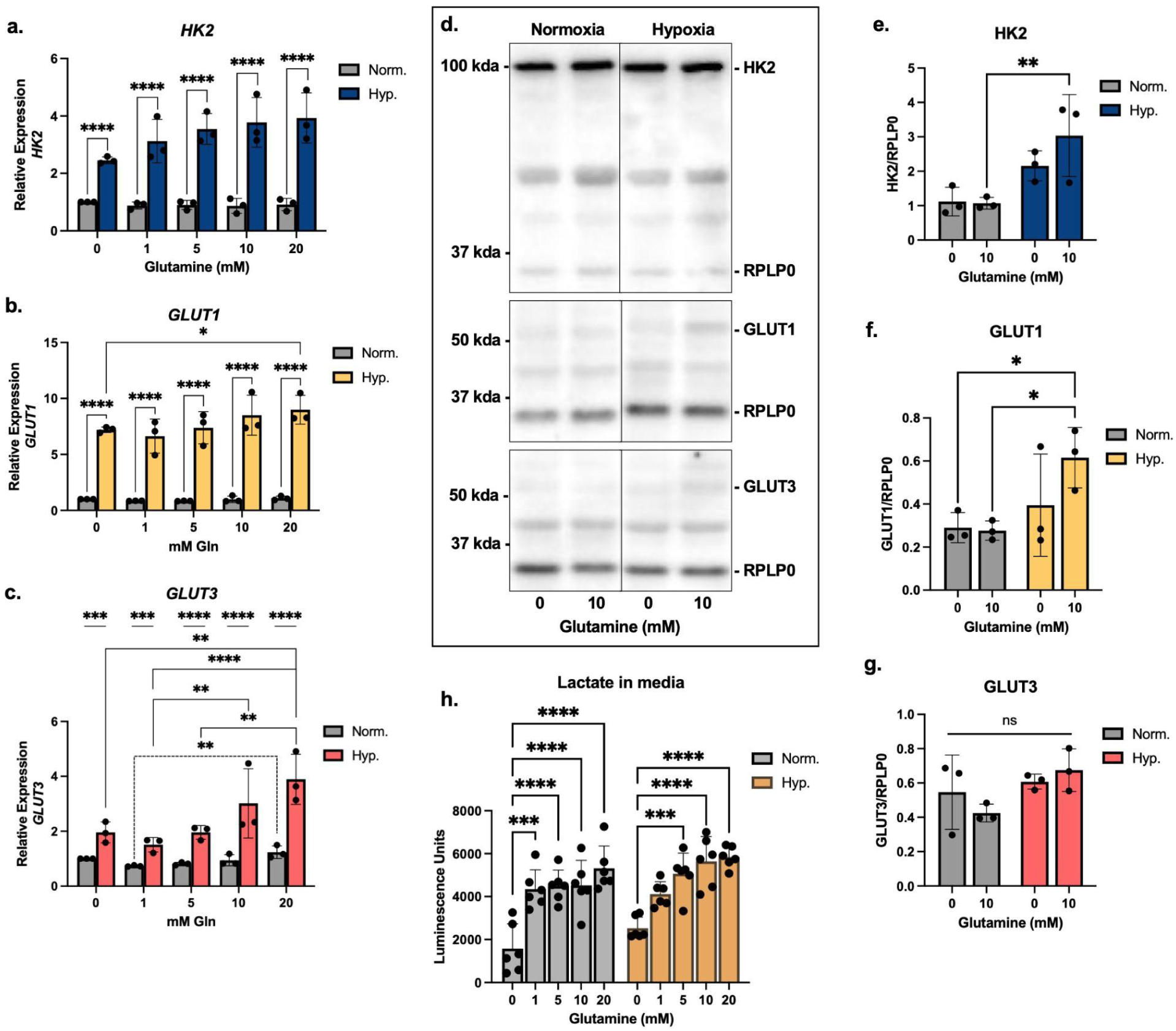
Glutamine enhances hypoxia-induced changes to glycolytic markers. LECs were supplemented with 0-20 mM of glutamine and then incubated in normoxic (∼21% O_2_; Norm.) or hypoxic (< 3% O_2_; Hyp.) conditions. Cells were harvested after 24 hours for gene and protein expression, while media was collected for metabolite analysis after 12 hours. Relative gene expression of **a)** *Hexokinase II (HK2)*, **b)** *Glucose Transporter 1 (GLUT1),* and **c)** *Glucose Transporter 3 (GLUT3)* was determined using the ΔΔCt method with *RPLP0* as an endogenous control. n = 3. Error bars represent the fold change standard deviation. Statistical significance was determined using delta Ct via two-way ANOVA with Tukey’s Test. **d)** Western blot of HK2, GLUT1, GLUT3, and RPLP0 after treatment with 0 mM (-) or 10 mM (+) of glutamine. **e-g)** Protein expression was quantified using ImageJ and normalized to RPLP0 . n = 3. **h)** Lactate in media was measured using the Promega Lactate-Glo assay, in which the amount of lactate is proportional to luminescence. n = 6. Statistical significance was determined via two-way ANOVA with Tukey’s Test. *P<0.05; **P<0.01; ***P<0.001; ****P<0.0001.

When examining protein expression using western blot (**Fig 2d)**, cells were supplemented with 10 mM (+) or 0 mM (-) of glutamine before normoxic or hypoxic incubation. With the exception of HK2, which was significantly increased by hypoxia in the presence of glutamine, changes in protein expression did not reach the level of statistical significance. However, hypoxic LECs supplemented with glutamine do show non-significant increases in HK2, GLUT1, and GLUT3 production compared to those deprived of glutamine **(Fig 2e-g)**. Lastly, the presence of glutamine significantly increased the presence of lactate in cell media in both normoxic and hypoxic conditions **(Fig 2h)**. Together, these results demonstrate a positive correlation between glutamine availability and glycolysis in hypoxic LECs.

#### 2.2.2. Lowering glutamine uptake decreases glycolytic gene expression and lactate production

To further define the relationship between glutamine availability and glycolysis, LECs were treated with V-9302, a competitive inhibitor of glutamine transport protein SLC1A5, also known as ASCT2 (Schulte et al., 2016; van Geldermalsen et al., 2016). It should be noted that SLC1A5 can also transport other neutral amino acids such as alanine, serine, and cysteine; however glutamine is its preferred substrate (Oppedisano et al., 2007; Scalise et al., 2020). Treatment with V-9302 significantly reduced glutamate production by hypoxic LECs, indicating a successful reduction of glutamine transport into the cell **(Fig 3a).** In hypoxic conditions, lowering glutamine transport also significantly decreased lactate production **(Fig 3b)**. V-9302 treatment also resulted in non-significant reductions in hypoxic *HK2* and *GLUT1* gene expression **(Fig 3c-d)**. Conversely, lowering glutamine transport significantly increased the hypoxic expression of *GLUT3*, supporting the idea that glucose transport is upregulated as compensation for decreased glutamine availability **(Fig 3e)**. These results suggest that, although hypoxia is the primary driver of LECs’ metabolic shift towards glycolysis, glutamine plays a supporting role in this change.

**Figure 3.**
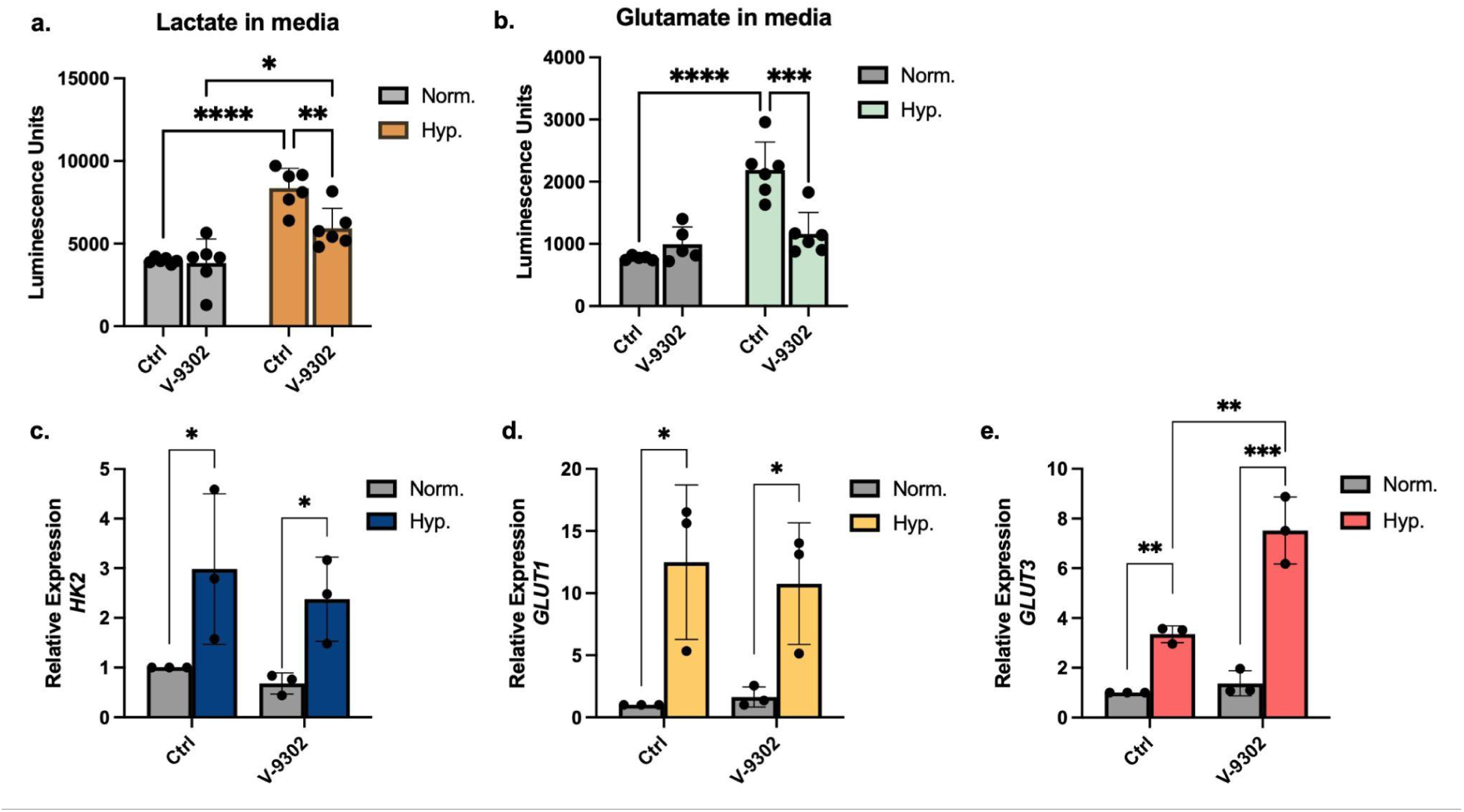
Blocking glutamine transport lowers glycolytic indicators but increases glucose transport gene expression in hypoxic conditions. LECs were treated with 10 μM of V-9302 for 48 hours prior to treatment. Before hypoxic incubation, fresh media containing V-9302 was provided. **a)** Glutamate in media as measured by Promega Glutamate-Glo assay at 12 hours, in which luminescence is proportional to the amount of glutamate in cell media. n = 5 for normoxic conditions, n = 6 for hypoxic conditions. **b)** Lactate in media as measured by Promega Lactate-Glo assay at 12 hours. Lactate is proportional to luminescence. n = 6. **c-d)** Relative gene expression of *Hexokinase II (HK2)*, *Glucose Transporter 1 (GLUT1), and Glucose Transporter 3 (GLUT3)* was determined using the ΔΔCt method, with *RPLP0* as the endogenous control. n=3. Error bars represent the fold change standard deviation. Statistical significance for all assays was determined using delta Ct via two-way ANOVA with Tukey’s Test. *P<0.05; **P<0.01; ***P<0.001; ****P<0.0001.

### 2.3. Glutamine causes functional changes in LECs

Having observed a positive relationship between glutamine and glycolysis in hypoxic LECs, we further tested how glutamine availability affected functions known to be powered by glycolytic activity, including proliferation, migration, and vessel formation (De Bock et al., 2013; Wu et al., 2021).

#### 2.3.1 Glutamine supports LEC proliferation and cell migration

To investigate the effect of glutamine supplementation on LEC proliferation, we measured DNA synthesis using a ThermoFisher Click-iT™ Plus EdU AlexaFluor™ 488 Imaging Kit. Cells which incorporated EdU were undergoing DNA synthesis (S-phase) at the time of fixing and displayed green fluorescence **(Fig 4a)**. Cell proliferation was then quantified by the percentage of cells in S-phase **(Fig 4b)**. We observed that the presence of glutamine significantly increased the number of normoxic LECs in S-phase, from under 2% in glutamine-free media to 16% with glutamine. In hypoxia, the presence of glutamine increased S-phase cells from approximately 1% to 3%. Hypoxia significantly reduced LEC proliferation, as such, glutamine appears to offer some protection against hypoxia-induced decreases in EC proliferation caused by oxidative stress (Hong et al., 2024).

**Figure 4.**
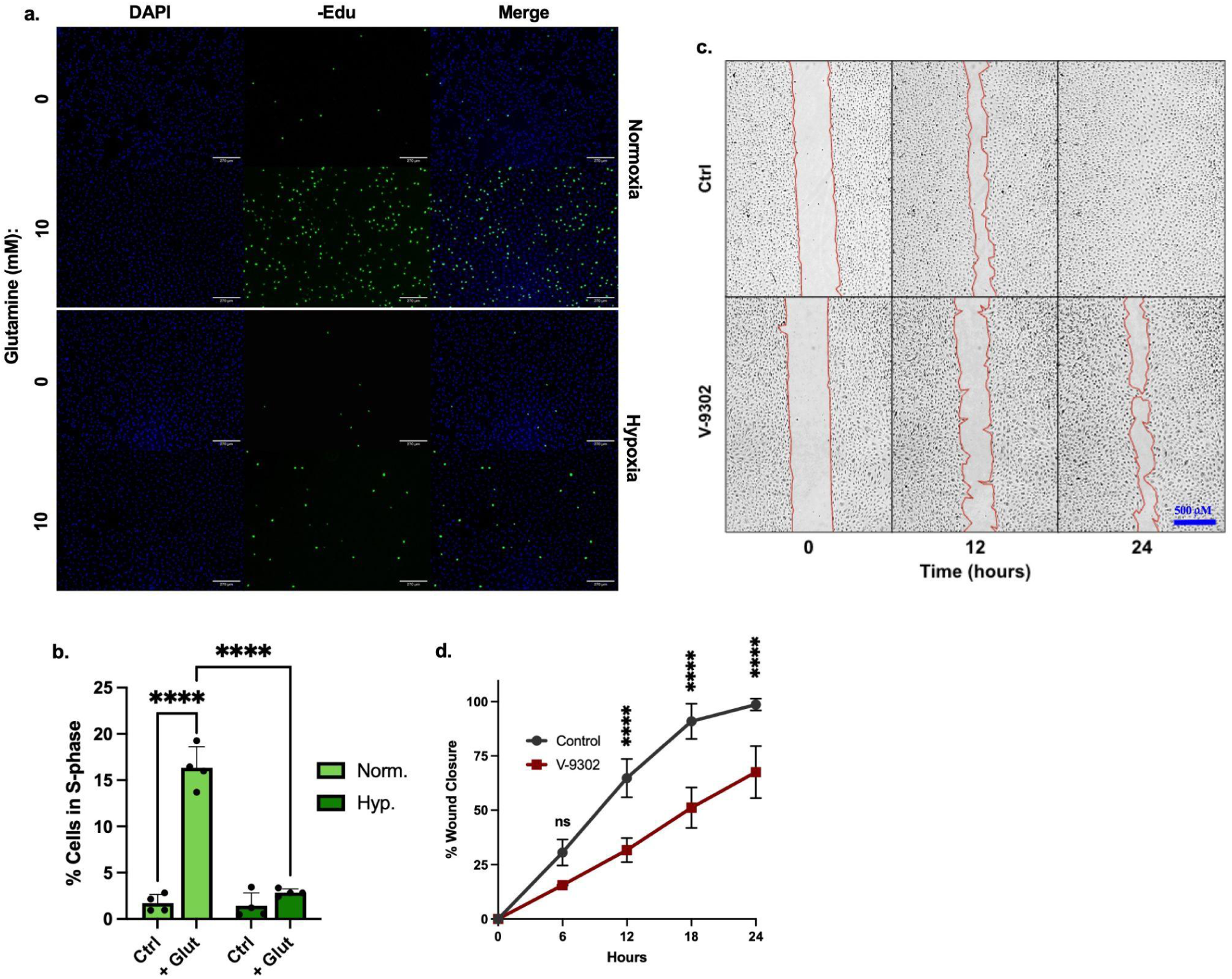
Glutamine increases LEC proliferation and migration. **a)** LEC proliferation after treatment with 0 mM (Ctrl) or 10 mM (+ Glutamine) in normoxic and hypoxic conditions. Cell nuclei were stained with DAPI (blue), while nuclei of cells in S-phase were stained green using an Invitrogen Click-It -Edu assay. Scale bar (white) = 270 μm. **b)** Cell proliferation as measured by percent cells in S-phase. n = 4. **c)** Representative images of wound healing assay in normoxic conditions at 0, 12, and 24 hours. Scale bar (blue) = 500 µm. **d)** Quantification of wound-healing assay as percent wound closure. n = 4 for control group; n = 3 for V-9302 group. Statistical significance for all quantifications determined using two-way ANOVA with Tukey’s Test. *P<0.05; **P<0.01; ***P<0.001; ****P<0.0001.

The impact of glutamine deprivation on LEC migration was observed using a wound-healing assay. LECs were seeded in Ibidi two well culture inserts and allowed to grow to confluency in either MV2 or MV2 containing 10 µM of V-9302 in normoxic conditions. Upon removal of the insert, the cells were imaged every 6 hours for 24 hours using an Agilent BioTek Lionheart Automated Microscope. Representative images at 0, 12, and 24 hours are shown in **Fig 4c**. The area of the wound was calculated using the manual segmentation option in the ImageJ Plugin Wound Healing Size Tool (Suarez-Arnedo et al., 2020). Untreated LECs showed significantly faster wound healing than V-9302 treated cells. While untreated LECs approached full wound closure by 24 hours, LECs treated with V-9302 approached 70% wound closure **(Fig 4d)**. This indicates that blocking glutamine transport slows the collective LEC migration needed for wound healing.

### 2.4. Impact of glutamine on LEC tube formation oxygen-dependent

To investigate the impact of glutamine supplementation and deprivation on *in vitro* lymphatic sprouting, a 2D tube formation assay was performed (Alderfer et al., 2021). LECs stained with Invitrogen CellTracker™ Red (Thermo Fisher Scientific) were seeded on Matrigel-coated Ibidi μ-slides at a density of 10,000 cells/well in supplemented MV2 which contained 10 mM glutamine, 15 mM glutamine, or 10 μM of V-9302. These concentrations were chosen as the LECs did not form vascular networks in glutamine-free DMEM, and basal MV2 media contains 10 mM of glutamine. The slides were then incubated for 6 hours in normoxic or hypoxic conditions. The vascular networks were then quantified with the ImageJ Plugin Kinetic Analysis of Vasculogenesis (Alderfer et al., 2021; Varberg et al., 2018). The average vessel length, number of closed loops, and tubes/nodes ratio were examined (Alderfer et al., 2024, 2021; Montes Pinzón et al., 2025) (**Fig 5 b-d**). Longer vessels and increased looping indicate more developed, inter-connected networks, while short vessels which form few loops indicate fragmentary vasculature (Jeong et al., 2023; Montes Pinzón et al., 2025). The tubes/nodes ratio can indicate both level of branching or fragmentation and relies on the other two metrics to provide context.

**Figure 5.**
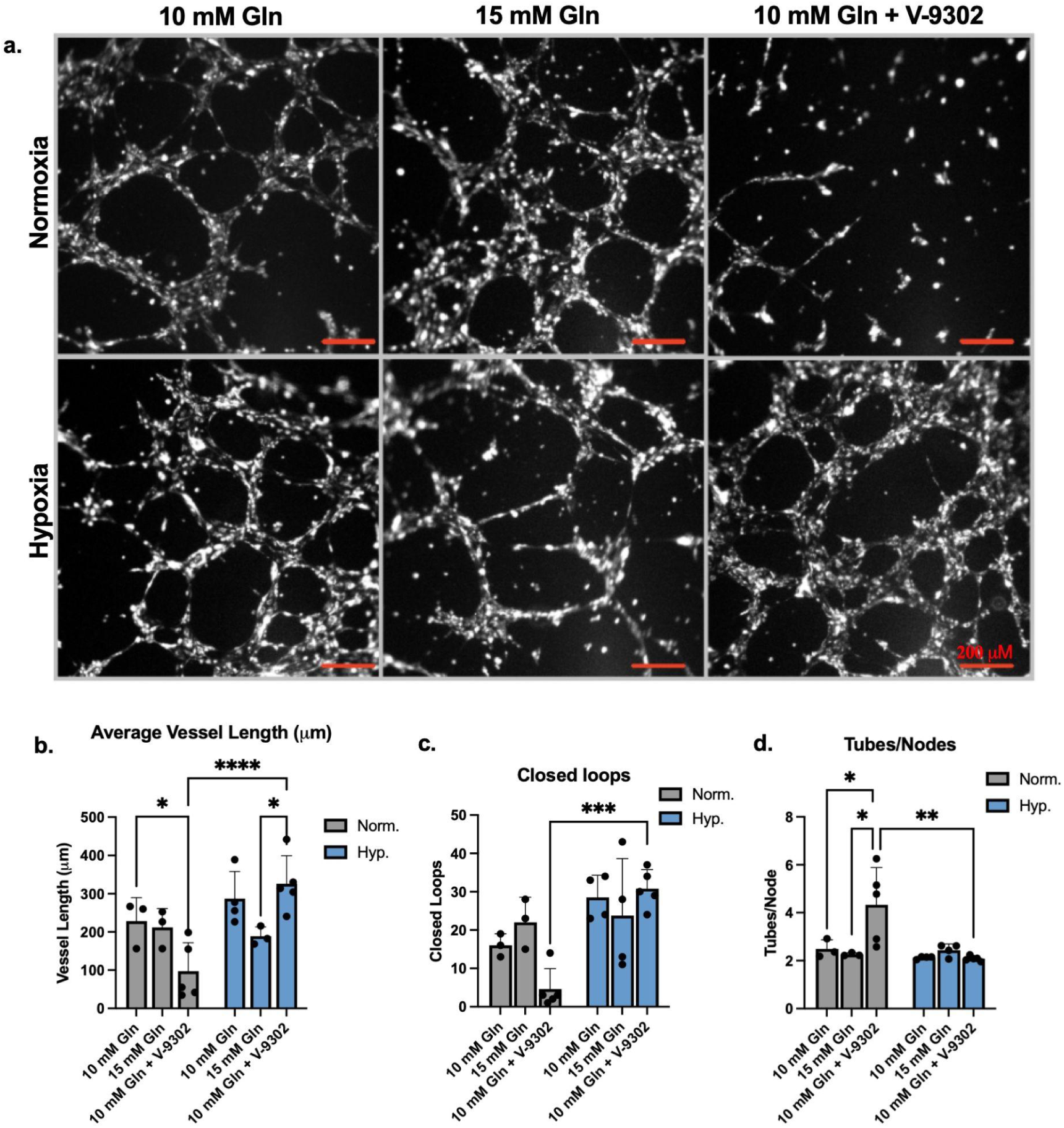
Impact of glutamine on LEC tube formation oxygen-dependent. LECs were stained with CellTracker Red and seeded on matrigel in MV2 (10 mM Gln; n=3 normoxia; n=4 hypoxia), MV2 + 5 mM glutamine (15 mM Gln; n=3 normoxia; n=4 hypoxia), or MV2 + 10 μm of V-9302 (10 mM Gln + V-9302; n=5 normoxia; n=5 hypoxia). The cells were then incubated in normoxic or hypoxic conditions for 6 hours. **a)** Representative images of 2D vascular networks. Scale bar (red) = 200 μm. **b-d)** Quantification of **b)** average vessel length, **c)** closed loops and **d)** tubes/nodes ratio in vascular networks using Image J Plugin Kinetic Analysis of Vasculogenesis. Statistical significance determined using two-way ANOVA with Tukey’s Test. *P<0.05; **P<0.01; ***P<0.001; ****P<0.0001.

We found that the effect of glutamine supplementation on lymphatic tube formation was dependent on oxygen. In normoxic culture, V-9302 treatment resulted in significant decreases in vessel length and increase in tubes/nodes ratio, as well as a non-significant decrease in closed loops formed, which matches the fragmentary network seen in **Fig 5a**. Meanwhile, LECs treated with 15 mM of glutamine had a slight increase in the number of closed loops (**Fig 5c**).

In hypoxic conditions, the opposite effect was observed. In comparison to cells treated with 10 mM glutamine or V-9302, LECs supplemented with additional glutamine (15 mM total) formed shorter vessels with fewer closed loops and higher tubes/nodes ratio (**Figure 5b-d**). LECs treated with V-9302, on the other hand, showed a morphology closer to that of the control group. This indicates that while glutamine availability can control vascular morphology, the effect is determined by environmental conditions.

## 3. Discussion

Glycolysis has a vast influence on lymphatic cell proliferation, migration, and lymphangiogenesis. This is especially true in hypoxic conditions, where glycolysis is further increased by both a lack of oxygen for cellular respiration and upregulation of glycolytic genes by HIF transcription factors. Therefore, small changes in glycolytic output can result in large changes to LEC behavior.

In this study, we investigated whether glutamine metabolism could influence glycolysis in hypoxic lymphatic endothelial cells. First, we showed that hypoxic conditions increase the amount of glutamate produced by LECs and therefore the amount of glutamine the cells are metabolizing. Next, we found that gene and protein expression of GLUT1, GLUT3, and HK2 increased with increasing glutamine concentrations under hypoxic conditions. LECs also produced more lactate when exposed to higher glutamine concentrations regardless of oxygen availability. The opposite effect was observed when glutamine transport was blocked using SLC1A5 inhibitor V-9302. It should be noted that lactate can also be a byproduct of glutaminolysis when glutamine is used for fatty acid synthesis, so changes in glycolysis may not be the only contributor to changes in LEC lactate production (DeBerardinis et al., 2007). Even so, the combined results indicate that glutamine availability influences glycolysis in hypoxic LECs.

Given that hypoxia is the main driver of metabolic shifts towards glycolysis, we then sought to determine if the changes to glycolysis caused by glutamine result in tangible differences in LEC behavior. For this, we examined LEC proliferation, migration, and vessel formation. We found that glutamine significantly increases LEC proliferation in normoxia and provides some protection against hypoxia-induced decreases in proliferation. This data agrees with previous studies on other endothelial cell types (Kim et al., 2017; Peyton et al., 2018; Wei et al., 2025). We also found that blocking glutamine transport with V-9302 slows LEC migration, which coincides with Peyton et al.’s (2018) observation that glutaminase-1 stimulates the migration of endothelial cells (Peyton et al., 2018). In a recent study comparing LEC and BEC metabolism, Durot et al. also found that inhibiting glutamate dehydrogenase over a 5-day period resulted in an LEC-specific decrease in cell migration, which was then rescued with dimethyl-α-ketoglutarate supplementation (Durot et al., 2025). It should be noted that Kim et al. (2017) did not report a link between glutamine and endothelial cell migration (Kim et al., 2017). This alternative finding could be attributed to differences in cell culture and treatments, and it points to the need to confirm these results *in vivo* in future studies.

Lastly, we examined the impact of glutamine supplementation and deprivation on lymphatic tube formation. While glutamine increased network formation in normoxic conditions, increasing glutamine availability in hypoxic conditions led to the generation of thinner tubes with less connectivity. This result was reversed with V-9302 treatment. We hypothesize that these changes in morphology are due to glutamine’s influence on the glycolytic activity of the cells. Hypoxia increases endothelial cell glycolysis through the HIF1α pathway, leading to the increased network formation seen in the control group. When glycolysis is further increased with glutamine supplementation, the rate of LEC migration increases as seen in **Fig 4c-d**, leading to thinner vessels. V-9302 attenuates any glutamine-induced increases to glycolysis, causing the vessel morphology to reflect the control hypoxia group.

While this study did not explore the mechanism by which glutamine availability influences hypoxic LEC glycolysis, previous research on glutamine metabolism offers several plausible mechanisms, including O-GlcNAcylation, mTORC1 signaling, and cellular response to redox homeostasis. O-GlcNAcylation is a post-translational addition of O-linked *N*-acetylglucosamine (O-GlcNAc) to Ser and Thr residues on various proteins. This molecule is made from glucose, glutamine, and acetyl-CoA through the hexosamine biosynthetic pathway (Yang and Qian, 2017). O-GlcNAcylation is increased by both increased nutrient availability and hypoxia. The O-GlcNAcylation of enzymes associated with glycolysis, such as G6PD, PGK1, PMK2, and PFKFB3, result in an increase of glycolytic activity (Cheng et al., 2024; Lei et al., 2020). Another possibility also involves nutrient sensing through the MTOR pathway. Work by Han et al. (2025) has uncovered that glycolysis and glutamine metabolism are both controlled by mTORC1 through Myc-HK2/GLS signaling during lymphatic development (Han et al., 2025). Other metabolic studies have shown that glutamine and glutaminolysis both activate mTORC1 (Durán et al., 2012; Meng et al., 2020; Tan et al., 2017). Another potential mechanism involves glutamine’s role in redox homeostasis. While excessive reactive oxygen species (ROS) can inhibit glycolysis, glutathione, a metabolite of glutamine, can scavenge ROS. Therefore, increased glutamine flux in hypoxic conditions may relieve ROS inhibition of glycolysis (Mullarky and Cantley, 2015). Future studies *in vivo* will be needed to confirm these relationships, as well as the ability of glutamine to influence glycolysis at concentrations that reflect those found in human plasma, approximately 0.4 - 0.9 mM (Bergström et al., 1974; Cruzat et al., 2018; Rodas et al., 2012).

This study identifies glutamine as a supporter of lymphatic endothelial cell glycolysis under hypoxic conditions **(Fig 6)**. It can be used as a tool to modulate lymphatic endothelial cell proliferation, migration, and vessel formation in normoxic and hypoxic conditions. This work provides evidence that indirect targeting of endothelial cell glycolysis is a viable method to control lymphangiogenesis. Potential applications include using glutamine to improve lymphatic vascularization in *in vitro* tissue models or spur wound healing in otherwise healthy tissues. Reducing glutamine uptake, on the other hand, could be used to normalize lymphatic vasculature in diseases with chronic hypoxia, such as solid malignancies or secondary lymphedema.

**Figure 6.**
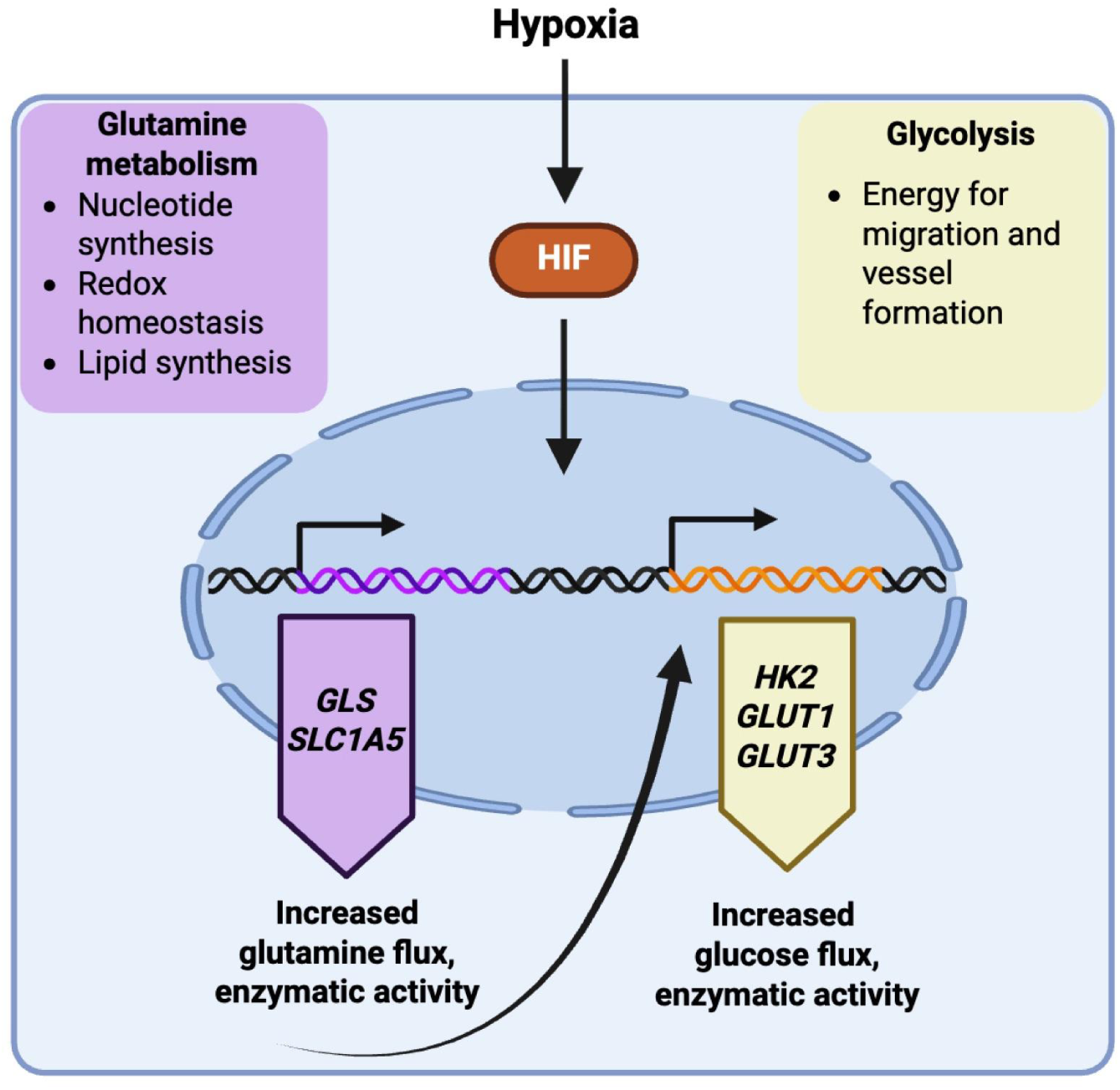
Summary of relationship between glutamine and glycolysis in hypoxic LECs. Hypoxia stabilizes HIF transcription factors, which increase the transcription of genes related to glutamine metabolism as well as glycolysis. Increased glutamine availability results in an increase in glycolytic activity. Image made in BioRender.

## 4. Materials and Methods

### 4.1 Cell Line and Culture

Juvenile Human Dermal Lymphatic Endothelial Cells (Promocell, C-11216) expanded in PromoCell Endothelial Growth Basal Medium MV2 with associated SupplementMix (C-22022) as previously described (Alderfer et al., 2024; Montes Pinzón et al., 2025; Saha et al., 2023). Unless stated otherwise, MV2 refers to complete media (MV2 Basal Medium + SupplementMix). Media was replaced every other day, and cells were passaged upon reaching above 80% confluency using the Promocell DetachKit (C-41220). Cells were used between passage 6 and passage 8 for experiments.

### 4.2 Cell Treatments

#### 4.2.1 Glutamine-supplemented media

To control glutamine concentrations, L-glutamine (Thermo Fisher Scientific, A14201.30) was dissolved in low-glucose, glutamine-free DMEM (Millipore Sigma, D5546) with or without MV2 SupplementMix to a concentration of 20 mM. It was then hand filtered through a 0.2 μm PES membrane before being diluted to reported concentrations. Prior to glutamine treatments, LECs were glutamine-starved by incubating for approximately 16 hours in glutamine-free DMEM with or without MV2 SupplementMix.

#### 4.2.2 Blocking SLC1A5

Glutamine transport inhibitor V-9302 (Millipore Sigma, SML3525) was dissolved in DMSO to a concentration of 3.71 mM and then added to MV2 to a final concentration of 10 μM. LECs were treated for 48 hours before being replaced with fresh media containing V-9302 for experiments. The concentration and duration of V-9302 treatment was based on similar *in vitro* treatments in the field of cancer research (Edwards et al., 2021; Schulte et al., 2018).

#### 4.2.3 Hypoxic Culture

For hypoxic treatments, the incubator was flooded with nitrogen in order to achieve a final gas ratio of 94% nitrogen, 5% carbon dioxide, and 1% oxygen. Oxygen levels in media were measured with PreSens Optical Oxygen Sensor (Regensburg, Germany) to confirm that oxygen levels in the media reached < 3% O_2_ (∼21 mmHg pO_2_). Although there is no universal standard for hypoxic treatment, 3% is considered moderate hypoxia (Vásquez Vélez et al., 2025). Normoxic cells were cultured at ambient oxygen (approximately 19-21% O_2_ or ∼140 mmHg pO_2_) in a separate incubator. Unless otherwise stated, cells were incubated in normoxic or hypoxic conditions for 24 hours. An overview of cell treatments is available in **Fig 7**.

**Figure 7.**
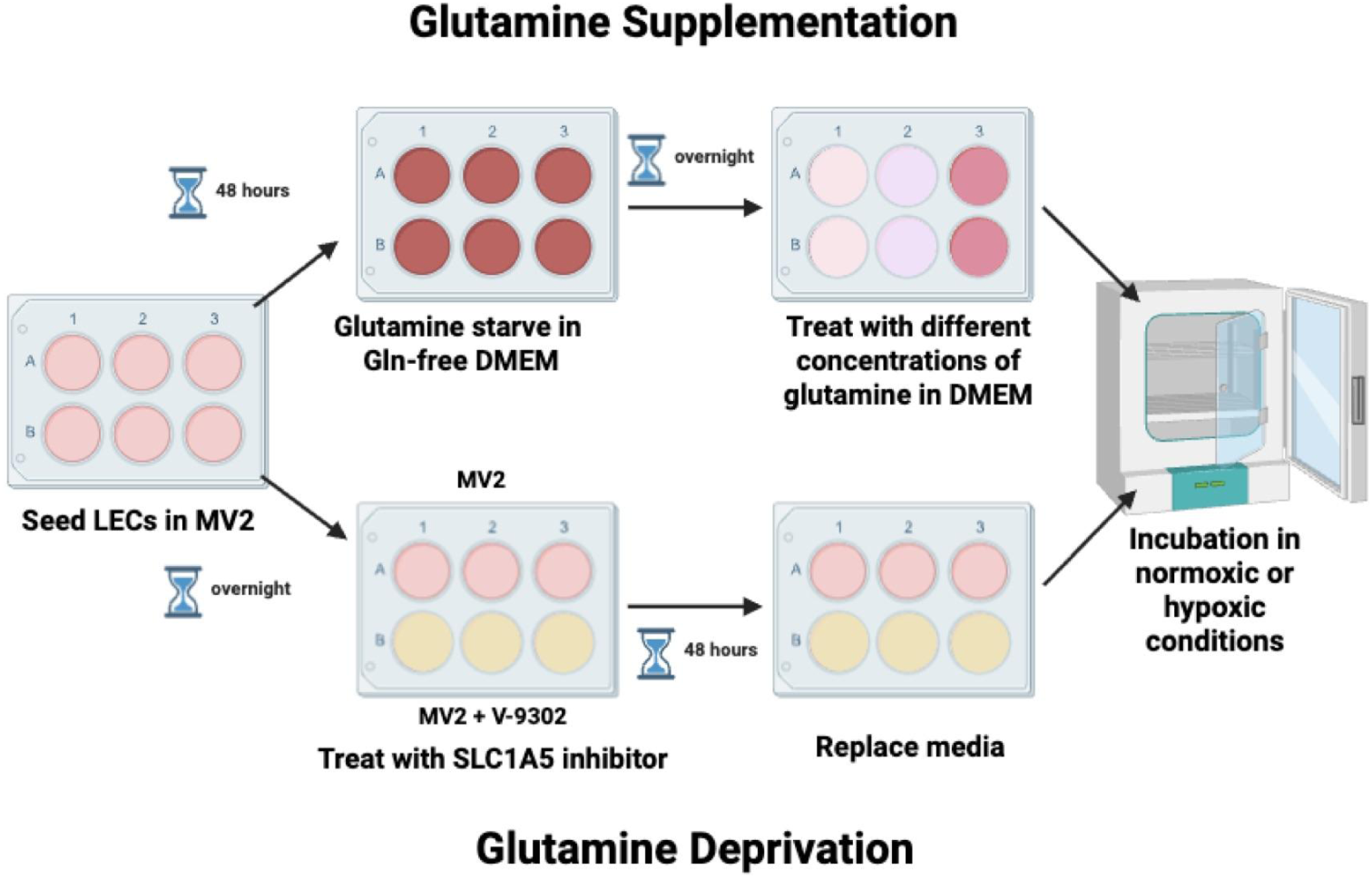
Schematic of cell treatments. Image made in BioRender.

### 4.3 HIF1A staining and quantification

20,000 cells in 0.5 mL of MV2 were seeded on top of 15 mm glass coverslips in a 24 well plate and incubated for approximately 48 hours. Afterwards, the media was replaced and cells were cultured for an additional 24 hours in normoxic or hypoxic conditions. Prior to staining, samples were fixed in 3.7% formaldehyde for 15 minutes, permeabilized with 0.1% Triton-X for 10 minutes, and blocked in 1% bovine serum albumin (BSA) in phosphate-buffered saline (PBS) for 1 hour. The cells were then stained with 1:200 Mouse anti-HIF1A (Enzo Life Sciences, BML-SA287-0100) for 90 minutes followed by 1:500 Donkey anti-mouse 647 (Abcam, AB150107) for 2 hours. Lastly, nuclei were stained with 300 nM DAPI for 3 minutes. Samples were mounted on glass slides and imaged at 10x magnification on an Echo Revolve 2 Confocal Microscope. Quantification of nuclear HIF1A was performed in Image J2 (Version 2.14.0) using a custom program which thresholds DAPI to create regions of interest (ROIs) for each nuclei (Hall, 2025). Afterwards, the fluorescent intensity of Hif1α is measured within each ROI. The individual fluorescent intensities for all nuclei were recorded across four wells for normoxic and hypoxic conditions. Images of all samples are available in **Fig S1.**

### 4.4 qPCR

80,000 cells in 2 mL MV2 were seeded in 6-well plates and allowed to attach overnight before undergoing treatments described in **4.2.1** and **4.2.2.** The cells were lysed with Invitrogen TRIzol™ Reagent (1559602) and frozen at -80℃. mRNA was isolated using a RNeasy Mini Kit (Qiagen, 74104). Samples were normalized to RNA concentration as measured by Thermo Scientific NanoDrop 2000c Spectrophotometer and converted to cDNA using an Invitrogen High Capacity cDNA Reverse Transcription Kit with Rnase inhibitor (4374967). qPCR was performed using the QuantStudio 5 Real-Time PCR system. Afterwards, relative gene expression was calculated using the delta-delta Ct method, with *RPLP0* expression used as a hypoxia-stable endogenous control (Bence et al., 2009; Lafont et al., 2008). Primers for *HK2*, *GLUT1*, and *GLUT3* were sourced from Thermo Fisher Scientific TaqMan™ assays (Hs00606086_m1, Hs00892681_m1, and Hs00359840_m1 respectively).

### 4.5 Lactate and glutamate production

LECs were seeded at a density of 10,000 cells/well in 96 well plates and allowed to attach for approximately 24 hours before being serum-starved overnight in glutamine free DMEM, MV2, or MV2 with V-9302. They were then incubated for 12 hours in normoxic or hypoxic conditions. Media was collected from each well and diluted 1:50 in PBS. Relative concentrations of glutamine and lactate in the media were then measured using Promega Lactate and Glutamate-Glo™ assays (J5022 and J7022) according to manufacturer instructions. Luminescence was read using a Tecan Spark plate reader, and final values were obtained after performing mean background subtraction. Wells that fell below the mean background or wells identified as outliers using Prism Robust regression and outlier removal (ROUT) analysis (Q = 1%) were removed from the data set (see supporting information **Table S10.2**).

### 4.6 Western Blot

LEC samples were lysed with 1:10 RIPA buffer with 1:200 protease inhibitor. To obtain whole-cell protein, samples were centrifuged at 16,000 xg for 20 minutes, after which the supernatant was collected. Protein concentration was determined using Pierce™ Dilution-Free™ Rapid Gold BCA Protein Assay (Thermo Scientific, A55862) . SDS-PAGE was performed using a 12.5% acrylamide gel with 10 μg protein/lane. The protein was then transferred to a nitrocellulose membrane. After blocking with 3% BSA, blots were incubated with primary antibodies for a combination of rabbit anti HK2 (Abcam, ab227198), GLUT1 (Abcam, ab12683), or GLUT3 (Thermo Scientific, PA5-99486) and RPLP0 (Abcam, ab154970) at 1:1000 dilution each. The blots were then washed in TBS and incubated with 1:5000 Goat anti-Rabbit IgG HRP (Abcam, ab6721). After washing with TBS, the blots were treated with Pierce™ ECL Western Blotting Substrate (Thermo Scientific, 32106) and imaged using an UVP ChemiDoc-It^2^® Imager and quantified using Image J2. The intensity of target bands were normalized to the intensity of RPLP0 bands. Uncropped blots are available in **Figure S2**.

### 4.7 Proliferation assay

40,000 cells in 0.5 mL of MV2 were seeded on 12 mm circular glass slides in 24 well plates and allowed to attach for approximately 24 hours. The cells were then serum and glutamine starved in glutamine-free DMEM overnight (approximately 16 hours) before media was replaced with supplemented DMEM containing 0 or 10 mM of glutamine. LECs were then incubated for an additional 24 hours in normoxic or hypoxic conditions before undergoing a proliferation assay using a Click-iT™ Plus EdU AlexaFluor™ 488 Imaging Kit (Thermo Fisher Scientific, C10637). Cell nuclei were stained with DAPI. Cells which incorporated Edu, and were therefore undergoing DNA synthesis (S-phase) at the time of fixing, displayed green fluorescence. Images were thresholded in ImageJ, and the percentage of cells in S-phase was calculated by dividing the number of nuclei displaying green fluorescence by the total number of nuclei.

### 4.8 Scratch assay

Ibidi 2 well cell culture inserts were placed in 24 well plates and seeded with 70µL of LECs at a density of 80,000 cells/mL in MV2 per side. An additional 0.5mL of MV2 was added outside the insert. On day 2, the media was replaced with MV2 (control) or MV2 containing 10µM of V-9302. On day 3, the media was replaced with serum-free MV2 (control) or serum-free MV2 + 10µM of V-9302 and allowed to incubate overnight. On day 4, the cell culture inserts were removed, and the cells were given fresh MV2 or MV2 containing V-9302. The cells were then placed in an Agilent BioTek Lionheart Automated Microscope, where they were kept at 37℃ and 5% CO_2_ and imaged every 6 hours for 24 hours. The area of the wound was calculated using the manual segmentation option in the ImageJ Plugin Wound Healing Size Tool (Suarez-Arnedo et al., 2020). The percent wound closure was determined by dividing the area of the wound at different time points by its initial area. Images of all samples, as well as areas selected for measurement, are available in **Fig S4**.

### 4.9 Vessel formation assay

The day prior to the assay, LECs treated according to section **2.2.1** and **2.2.2** were stained with 3 μM of Invitrogen CellTracker™ Red (Thermo Fisher Scientific) for 30 minutes and then passaged. The day of, 10 μL of Matrigel was added to Ibidi 15 well μ-slides (81501) and cured for 2 hours at 37℃. Then, the slides were seeded with 10,000 LECs per well in 50 μL of MV2, MV2 supplemented with 5 mM glutamine, or MV2 with 10 µM of V-9302 and placed in normoxic or hypoxic incubators for 6 hours to form vessels. Images were taken at 4x magnification using an Echo Revolve 2 Confocal Microscope. Before quantification, the images were processed in Image J2 by applying the built-in Gaussian blur filter with a sigma of 5 and then manually adjusting brightness and contrast (see **Figure S5**). The processed images were then analyzed using the ImageJ Plugin Kinetic Analysis of Vasculogenesis (KAV) to quantify tube/node ratio, vessel length, and number of closed loops (Bui et al., 2022; Varberg et al., 2018).

### 4.10 Supplemental materials

Supplemental materials include images of all samples for HIF1A staining (Fig S1), proliferation assay (Fig S3), and wound healing assay (Fig S4). Uncropped western blot images are available in Fig S2, and raw and processed images from all samples are available in Fig S5. A consolidated list of reagents is available in Table S1. Raw data from all assays can be found in Table S2-S16.

Supplemental materials are available at: https://figshare.com/articles/preprint/Supporting_Information_b_i_Glutamine_Availability_Impact s_Lymphatic_Endothelial_Cell_Glycolysis_and_Lymphangiogenesis_in_Hypoxic_Environments_i_b_/30913952?file=60823957.

## Acknowledgments

We acknowledge support from the University of Notre Dame through “Advancing Our Vision” Initiative in Stem Cell Research, Harper Cancer Research Institute – American Cancer Society Institutional Research Grant (IRG-17-182-04), American Heart Association through Career Development Award (19-CDA-34630012 to D.H.-P.), National Science Foundation (2047903 and 2225601 to D.H-P.), and the National Institutes of Health (1R35-GM-143055 to D.H.-P.).

## Conflicts of interest

The authors declare no conflicts of interest

## Author Contributions

E.J., D.H.P., E.H. and M.S. conceived and planned the experiments. E.J. carried out experiments, conducted data analysis, and wrote the manuscript. E.H. designed the ImageJ code to quantify fluorescence in the nucleus of individual cells. T.H. and K.P. assisted in conducting experiments. All authors contributed to data interpretation and editing the manuscript.

## Notes

https://doi.org/10.6084/m9.figshare.30913952

### Competing Interest Statement

The authors have declared no competing interest.

### Summary of Updates

Paper has been re-formatted for clearer reading, with updated figures and an expanded summary section

