## Supporting Information Table 1 for "Blocking glutamine transport normalizes lymphatic vessels in hypoxic environments by attenuating glycolysis" (from another version of this preprint: version 2, 2026-02-17)

**Table S1.** List of materials and reagents.

| Assay | Product | Manufacturer | Catalog number |
|---|---|---|---|
| **Oxygen Monitoring** | SDR SensorDish™ Reader Set | PreSens; Regensburg, Germany | N/A |
| | OxoHydroDish™ 6 well plate | PreSens | N/A |
| **Cell Culture and Treatments** | Juvenile Human Dermal Lymphatic Endothelial Cells | Promocell; Heidelberg, Germany | C-11216, lot 483Z001.2 |
| | Endothelial Growth Medium MV2 + SupplementMix | Promocell | C-22022 |
| | DMEM low glucose, without L-glutamine | Millipore Sigma; Burlington, MA, USA | D5546 |
| | DetachKit | Promocell | C-41220 |
| | L-glutamine, 99% | Thermo Fisher Scientific; Waltham, MA, USA | A14201.30 |
| | V-9302 | Millipore Sigma | SML3525-5MG |
| **Immunostaining** | HIF-1α monoclonal antibody (H1α67) | Enzo Life Sciences; Farmingdale, NY, USA | BML-SA287-0100 |
| | Donkey Anti-Mouse IgG H&L (Alexa Fluor® 647) | Abcam; Cambridge; UK | AB150107 |
| | Invitrogen CellTracker™ Red CMTPX | Thermo Fisher Scientific; Waltham, MA, USA | C34552 |
| **PCR** | *Hexokinase II (HK2)* primer | Thermo Fisher Scientific | Hs00606086_m1 |

| | | | |
|---|---|---|---|
| | *Solute carrier family 2 member 1* (SLC2A1/*GLUT1*) primer | Thermo Fisher Scientific | Hs00892681_m1 |
| | *Solute carrier family 2 member 3 (SLC2A3/GLUT3)* primer | Thermo Fisher Scientific | Hs00359840_m1 |
| | Ribosomal protein lateral stalk subunit P0 (RPLP0) primer | Thermo Fisher Scientific | Hs00420895_gH |
| **Metabolite Assays** | Lactate-Glo™ | Promega; Madison, WI, USA | J5022 |
| | Glutamate-Glo™ | Promega | J7022 |
| **Western Blot** | HK2 antibody | Abcam | ab227198 |
| | GLUT1 antibody | Abcam | ab14683 |
| | GLUT3 Polyclonal Antibody | Thermo Fisher Scientific | PA5-99486 |
| **Proliferation** | Click-iT™ Plus EdU AlexaFluor™ 488 Imaging Kit | Thermo Fisher Scientific | C10637 |
